# Plasmodium falciparum aquaglyceroporin (PfAQP) orchestrates glycerol-dependent lipid metabolism, membrane biogenesis, metabolomic flux, parasite growth and stress adaptation

**DOI:** 10.64898/2025.12.23.696167

**Authors:** Monika Narwal, Md Muzahidul Islam, Madiha Abbas, Yoshiki Yamaryo-Botté, Pawan Malhotra, Zeeshan Fatima, Cyrille Y. Botté, Asif Mohmmed

## Abstract

Aquaglyceroporins (AQPs) are integral membrane channel proteins that facilitate transport of water and other solutes such as glycerol, thereby playing vital roles in cellular homeostasis. Here, we functionally characterized *Plasmodium falciparum* aquaglyceroporin (*Pf*AQP) through localization, inducible knockdown, and comprehensive metabolomic analyses revealing its key role in lipid homeostasis and metabolomic flux. *Pf*AQP localizes to the parasite plasma membrane and food vacuole membrane, at critical interfaces for nutrient and metabolite exchange. A *glm*S-ribozyme mediated inducible knock-down of *Pf*AQP severely impairs parasite development from trophozoite to schizont, leading to marked suppression in parasite growth. Ultra-Expansion Microscopy (UExM) analyses show profound defects in membrane biogenesis, along with disruption in cytokinesis under *Pf*AQP knock-down conditions. Detailed lipidomic analyses suggests that *Pf*AQP disruption cause altered lipid homeostasis, characterized by reduced membrane phospholipid biosynthesis, impaired lipid scavenging, and accumulation of storage lipids. Metabolomic analyses further indicate metabolic dysregulation characterized by reduction in amino acid levels and dysregulated mitochondrial metabolism, including impaired tricarboxylic acid (TCA) cycle. Strikingly, *Pf*AQP expression gets transiently upregulated under nutrient starvation condition, and its ablation drastically compromise parasite recovery, highlighting its role in adaptive responses. Overall, these results establish *Pf*AQP as a critical regulator of glycerol-dependent metabolic pathways, orchestrating membrane biogenesis, energy metabolism, and stress resilience—functions indispensable for parasite growth and survival.

## Introduction

Malaria exerts an enormous toll on global public health especially in developing world. The World Malaria Report of 2024 reports that there were an estimated 263 million malaria cases worldwide ^1^. The human malaria parasite *Plasmodium falciparum* is responsible for most malaria-related mortality. Emergence of resistance to artemisinin and partner drugs in *P. falciparum* has necessitated the development new antimalarials. Identification of unique and essential metabolic pathways of the parasite is a pre-requisite for development of novel drug candidates.

*P. falciparum* is a dynamic organism that carries out its life cycle in two different hosts. Inside the human host, it invades hepatocytes and red blood cells and is able to differentiate into distinct morphological stages during its life cycle. As soon as free parasites invaded an erythrocyte, a series of major transformations begin accompanied by rapid growth from progression of ring to schizonts. This 48-h asexual blood stage development produces up to 16-25 daughter cells called merozoites, which in turns leads to biosynthesis of enormous amount of membrane lipids requirement ^2, 3^. We have recently shown that he parasite uses a lysophospholipase (LPL3) at the host-parasite interface to generate fatty acids from host acquired phospholipids, these fatty acids are stored and subsequently mobilized for synthesis of phospholipids essentially required for membrane biogenesis or daughter merozoites ^4^. Due to the absence of a functional cholesterol biosynthesis pathway, parasite membranes predominantly comprise glycerolipids-primarily glycerophospholipids-built upon a glycerol backbone ^5, 6^. In *Plasmodium*, the glycolysis is a major provider of the glycerolipid backbone, glycerol-3-phosphate (G3P), however, the glycerol pool has been also been suggested to be a source for the same ^5, 7^. Unlike many prokaryotes and some eukaryotes, *Plasmodium* parasites appear to lack a complete enzymatic pathway for the de novo synthesis of glycerol from carbohydrate precursors via glycerol-3-phosphate phosphatase. Instead, glycerol-3-phosphate (G3P), the central precursor for glycerolipid synthesis, is primarily generated through phosphorylation of imported glycerol by glycerol kinase (GK) or via reduction of dihydroxyacetone phosphate (DHAP) by G3P dehydrogenase (G3PDH). DHAP itself arises from glycolysis, allowing the parasite to partially bypass direct glycerol uptake under certain metabolic conditions. However, extracellular glycerol acquisition is also critical, especially when lipid biosynthetic demand is high, such as during schizogony.

In higher organisms, the aquaporins (AQP) family proteins are thought to be one of the major players in the glycerolipid metabolism ^8^. AQPs belong to the major intrinsic protein (MIP) family, a superfamily of transmembrane protein channels and exist in almost all living organisms including animals, plants, bacteria and viruses ^9^ consisting of 6 α-helical transmembrane domains (TM) and five connecting loops ^10, 11^. Based on permeability, AQPs are categorized into water specific channels (orthodox aquaporins) and aquaglyceroporins, which are permeated by glycerol and some other solutes. Orthodox aquaporins serve vital functions in the fluid/water homeostasis ^12^, whereas aquaglyceroporins are involved in metabolism physiological processes like triglycerides metabolism, specially glycerolipid biosynthesis, and TCA cycle ^13^. Several water permeable porins like AQP0, AQP1, AQP2, AQP4, AQP5, AQP6 and AQP8 have been identified across eukaryotes and prokaryotes. On the other hands, AQP3, AQP7 and AQP9 are known to be permeable to glycerol and urea, ammonia or other small, uncharged molecules. This has also been established that these family of proteins play an important role in fasting or starvation state ^3^.

A single aquaglyceroporin is present in *P. falciparum* (*Pf*AQP), which is suggested to prevent the parasite from intoxication by facilitation the export of residual by-products ^14, 15^. However, the exact functional significance of *Pf*AQP and its role in parasite cellular homeostasis has not analysed. *Pf*AQP, is a member of aquaglyceroporin subfamily with the highest similarity to *Escherichia coli* glycerol facilitator (glpF) ^16^. Unlike its close structural homolog from *E. coli* (GlpF), *Pf*AQP not only conducts glycerol transportation but also water at high rates ^17, 18^. Sequence comparison shows almost identical residues in both filter regions of *Pf*AQP and GlpF. Although *Pf*AQP and GlpF share the same set of residues (Arg, Trp, and Phe) in their ar/R constriction, the NPA motifs of *Pf*AQP are subtly altered to Asn-Leu-Ala (NLA) and Asn-Pro-Ser (NPS). However, the water and glycerol permeability of *Pf*AQP were not affected by the mutation of the two varying amino acids back to canonical NPA motifs ^16^. Similarly, the aquaglyceroporin homologue in *Plasmodium berghei* (*Pb*AQP) is known to provides the major route for glycerol uptake into the parasite ^19^.

The role of host aquaporins is shown to play role during *Plasmodium* infection; the human AQP3 is found to be localized to the parasitophorous vacuole of red blood cells infected with *P. falciparum* ^20^. Further, the expression of AQP3 gets induced in human hepatocytes in response to *P. berghei* infection, which localizes to the PVM and contribute to the rapid replication and expansion of the parasite ^19^. Genetic disruption of AQP3 leads to a significant decrease in the parasite load of the *P. berghei* at liver stage.

As mentioned above, the *P. falciparum* genome encodes one aquaglyceroporin (*Pf*AQP Gene ID: PF3D7_1132800). The current work involves the functional characterisation of *P. falciparum* aquaglyceroporin (*Pf*AQP) to show its importance in parasite growth and survival, lipid homeoatasis and parasite energy metabolism. Our studies demonstrate that the *Pf*AQP is essential for parasite blood stage growth, and this effect on parasite growth was associated with a significant impairment phosoplipid biosynthesis as well as functionality of TCA cycle mostly due to reduced glycerol transport. This study highlights the importance of a novel metabolite uptake pathway in the parasite which is crucial for cellular homeostasis and parasite survival.

## Materials and methods

### Parasite culture, plasmid constructs, and parasite transfection

*P. falciparum* parasites, wild-type 3D7 strain and transgenic parasite lines, were cultured in 4% hematocrit in RPMI 1640 supplemented with 27 mM sodium bicarbonate, hypoxanthine (0.001%), and 0.5% Albumax I (Invitrogen) following the standard culture protocols ^21^. To generate GFP targeting plasmid construct, the entire open reading frame of the *Pf*AQP gene, (Gene ID: PF3D7_1132800) without stop codon (1–774 bp of mRNA sequence) was amplified using specific primers 1343A and 1344Aand was cloned into pSSPF2 vector ^22^ using *Bgl*II and *Avr*II restriction enzyme. Synchronized ring stage *P. falciparum* 3D7 parasite was transfected with 100 μg of purified plasmid DNA by electroporation (310 V, 950 μF) ^21^. The transfected parasites were selected over Blasticidin S drug (2.5 μg/ml). To generate the plasmid construct PfAQP-HA-T2A-WR-glmS, a C-terminal fragment of 600bp of sequence upstream of the stop codon of *pfaqp* was amplified using specific primers 1589A and 1590A from *P. falciparum* 3D7 genomic DNA and cloned in to *Kpn*I and *Xho*I sites of pHWGB vector. The transfected parasites were selected over 2.5 μg/ml of Blasticidin S (Gibco, Thermo Scientific). The parasites survived were subjected to selection using selection linked resistance by WR99210 drug to enrich parasites population where plasmid was integrated in the main genome. Integration of HA-*glm*S at the C-terminus was confirmed by PCR using specific primers (Supplementary Table S1), as well as by western blot analysis using an anti-HA antibody.

### Starvation Assay and Growth Recovery Assay

*P. falciparum* 3D7 parasites at ring stage were grown in complete RPMI or isoleucine-free RPMI for various increments of time (24 hpi, 30hpi and 34hpi). For growth recovery assay, the starved cultures were supplemented with isoleucine after starvation for various periods of time and were allowed to recover for an additional 72hr. Total parasitemia was assessed by counting new ring stage parasite.

### Parasite fractionation and Western blotting

Western blot analysis was used to evaluate the expression of fusion protein in transgenic *P. falciparum* parasite strains. In short, erythrocytes were lysed with 0.15% saponin to extract parasites from mixed stage culture. The parasite pellets were then suspended in Laemmli buffer, boiled, centrifuged, and the resulting supernatant was separated on 12% SDS-PAGE. After being separated from the gel, the fractionated proteins were placed onto the PVDF membrane (Merck Millipore) and blocked for two to three hours using blocking solution (1×PBS, 0.1% Tween-20, and 5% milk powder). After washing, the blot was incubated for three hours with primary antibodies diluted in dilution buffer (1×PBS, 0.1% Tween-20, and 2% milk powder): mouse anti-GFP (Roche, 1:1000); rabbit anti-BiP (1:10000) ^22^; and rat anti-HA (Roche, 1:1000). Claritymax ECL Substrate detection kit (BioRad) was used to visualize the bands after the blot had been washed and incubated for one hour with the respective secondary antibody (anti-rabbit, anti-rat, or anti-mouse, 1:20000) conjugated to HRP.

### Conditional knock-down and in-vitro growth assays analysis

For *glm*S-mediated conditional knock-down of target gene, tightly synchronized *Pf*AQP HA-*glm*S transgenic parasite cultures at the early ring stage (6–8 hpi) were incubated with glucosamine (2.5 mM) or solvent alone, and allowed to grow for three consecutive cycles. To assess the effect on the growth and morphology of the parasite, thin smears of parasite culture were made from each well at different time points (0, 24, 48, 72, 96, and 120 h) and stained with Giemsa stain for microscopic analysis. The parasitemia was determined in each cycle by counting the parasites in Giemsa-stained smears. The numbers of ring/trophozoite stage parasites per 1000 RBCs were determined in Giemsa-stained smears and percentage parasitemia was calculated to assess the parasite growth inhibition. Each assay was performed three times separately on different days.

### Organelle labelling, immuno-labelling, fluorescence microscopy, and Ultrastructure expansion microscopy

For indirect immunofluorescence assay, infected RBCs were fixed with a fixative solution, 4% w/v paraformaldehyde (Sigma) permeabilised with 0.1% v/v PBS-Triton X-100 and then blocked with 5% FBS-PBS. For immune-labelling, fixed culture was incubated with primary antibodies, Rat anti-HA (1:250) (Roche), Mouse anti-SERA5 (1:1000) followed by appropriate secondary antibody incubation goat anti-rat Alexafluor 488 (1:500), goat anti-rabbit Alexafluor 594 (1:500) or goat anti-mice Alexafluor 594 (1:500) (Invitrogen). Membrane structures of parasitized erythrocytes were labelled using 1 μM BODIPY™ TR Ceramide (Invitrogen). Nucleus was stained with DAPI (2 μg/ ml, Sigma Aldrich). Images were acquired using Nikon A1 confocal laser scanning microscope and analysed with Nikon NIS-Elements software (version 4.1).

U-ExM was performed as previously described with minor modification ^23, 24, 25^. Briefly, synchronized *Pf*AQP (control and iKD) schizonts stage parasites were percoll purified and were placed on poly-D-lysine-coated coverslips for 1hour at 37°C to settle in one well of a 24-well plate. Parasite cultures were set to 0.5% hematocrit, and 1 mL of parasite culture was added to the well containing the coverslip for 15 min at 37°C. Culture supernatants were removed and then fixed with 4% PFA for 20 minutes at 37°C and were washed three times with PBS. Following fixation, the coverslips were incubated with formaldehyde/acrylamide (FA/AA) overnight at 37°C. The next morning, to polymerize the gel, N,N,N’,N’-tetramethylethylenediamine (TEMED) and ammonium persulfate (APS) were swiftly added to monomer solution (19% w/w sodium acrylate [Sigma,], 10% v/v acrylamide [Sigma], 0.1% v/v N,N’-methylenebisacrylamide [Sigma] in PBS). Prior to gelation, FA/AA solution was removed from coverslips, and they were washed once in PBS. For gelation, 5 µL of 10% v/v tetraethylenediamine (TEMED; Thermo Fisher, Cat# 17919) and 5 µL of 10% w/v ammonium persulfate (APS; Thermo Fisher, Cat# 17874) were added to 90 µL of monomer solution and briefly vortexed. Subsequently, 35 µL was pipetted onto parafilm and coverslips were placed (cell side down) on top. Gels were incubated at 37°C for 30 min before being transferred to wells of a 6-well plate containing denaturation buffer (200 mM sodium dodecyl sulfate [SDS], 200 mM NaCl, 50 mM Tris, pH 9). Gels were incubated in denaturation buffer with shaking for 15 min, before separated gels were transferred to 1.5 mL tubes containing denaturation buffer. 1.5 mL tubes were incubated at 95°C for 90 min. Following denaturation, gels were transferred to 10 cm Petri dishes containing 25 mL of MilliQ water for the first round of expansion and placed onto a shaker for 30 min three times, changing water in between. Gels were subsequently shrunk with two 15 min washes in 25 mL of 1× PBS, before being transferred to 6-well plates for 30 min of blocking in 3% BSA-PBS at room temperature. After blocking, gels were incubated with primary antibodies, diluted in 3% BSA-PBS, overnight. Primary antibodies were used at the following dilutions: rat anti-HA (1:250) [Sigma], rabbit anti-*Pf*MSP1 (1:500), *Pf*CRT (1:250). The next day, gels were washed three times with 2 mL 0.5% Tween20 in PBS for 10 minutes at room temperature with agitation. Gels were then incubated in 1 mL of PBS with secondary antibodies including NHS-Ester (AlexaFluor 405, 1:100) protected from light, with agitation at room temperature. After 2.5 hours of secondary antibody incubation, gels were washed three more times with 2 mL 1% PBS + 0.5% Tween20 as before, then placed in a 10-cm dish filled with ddH2O. Water was replaced after 30 minutes, twice, and gels were allowed to expand overnight before imaging on a Nikon A1 confocal laser scanning microscope with a 100× magnification objective and a numerical aperture of 1.4. For gels stained with BODIPY TRc (Invitrogen), the fully expanded gel was incubated overnight at room temperature in 0.2% w/v propyl gallate (Sigma) solution containing BODIPY TRc (2 µM final concentration). For most of the displayed images, z-stack acquired on Nikon A1 confocal laser scanning microscope were processed in Fiji/ImageJ V1.54 software ^26^.

### Isolation of RNA, and quantitative RT-PCR

Parasites were harvested by saponin lysis and total RNA was prepared by the TRIzol based extraction following manufacturer’s protocol. Total RNA was reverse transcribed using the iScript cDNA Synthesis Kit (Bio-Rad) following manufacturer’s recommendations. Real-time amplification reactions were performed in triplicate on the Step OnePlus™ Real-Time PCR System (Applied Biosystems, Foster City, CA, USA) using a SYBR Green master mix from Biorad (CA, USA). Each reaction comprised equal amount of cDNA, 100 ng of both the gene-specific primers for *pfaqp* (1591A and 1592A), *Pf*18sRNA (18sF and 18sR), and 1x SYBR Green PCR mix. The threshold cycles (Ct values) generated by qPCR system were used to calculate fold change for relative quantitative analysis as described earlier ^27^.

### Lipidomic analysis

Lipidomics analysis was performed on four independent cell harvests of the control and iKD sets at 36 hpi. Highly synchronous parasite cultures were harvested and metabolically quenched via rapid cooling to 0°C by suspending the tube over a dry ice/100% ethanol slurry mix. The conventional Modified Bligh and Dyer method ^28, 29^ of total lipid extraction was utilised, involving the mixture of dichloromethane, methanol and water in a 2:2:1 v/v ratio. Lipid mixtures with ceramide and splash mix were employed as internal standards. Further, for the LC-MS run, the lipids were resuspended in 100 μL of ethanol and injected as described earlier ^30, 31^. Lipid chromatography was performed using the Exion LC system and a Waters AQUITY UPLC BEH HILIC XBridge Amide column (3.5 μm, 4.6 × 150 mm) with a single injection of 10 μL of sample. Buffer A (95% acetonitrile, 10 mM ammonium acetate, pH 8.2) and Buffer B (50%) were used to separate lipids, as described earlier ^32^. The Sciex QTrap 6500 triple quadrupole mass spectrometer was used for mass-spectrometry in a lower mass range. The probe’s parameters included curtain gas at 35 psi, temperature at 500°, source gas (SG1) and source gas (SG2) at 50 and 60 psi, and ionization voltage (IS) positive and negative mode at 5500 and 4500, respectively. The target scan period was 0.5 s, with 10 Da/s speed, 50,000 ms settling time, and 50,070 ms MR pause, as described earlier ^32^.

For total fatty analysis, total lipid was then derivatized to give fatty acid methyl ester (FAME) using trimethylsulfonium hydroxide (TMSH, Machenery Nagel) for total glycerolipid content. Resultant FAMEs were then analyzed by GC-MS as previously described ^33^. All FAME were identified by comparison of retention time and mass spectra from GC-MS with authentic chemical standards. The concentration of FAMEs was quantified after initial normalization to different internal standards and finally to parasite number.

### Metabolite extraction and relative quantitation

Metabolites were extracted using biphasic methanol-chloroform-water extraction protocol as described previously ^34^. Briefly, synchronized parasite cultures from both sets (control and - iKD) were collected by saponin lysis (0.05% in PBS). The parasite pellets were flash-frozen in liquid nitrogen and kept at –80 °C until extraction. Pellets were resuspended in ice-cold 80% methanol, sonicated for 10 minutes, then extracted with chloroform and water at a solvent ratio of 4:3:2 (methanol:chloroform:water, v/v/v). The upper aqueous phase was dried under vacuum using a SpeedVac concentrator (Thermo Scientific). TCA cycle and amino acid standards were added to the dry pellet, concentrated with methanol, and dried. Samples were resuspended in 20 μl of methoxyamine-HCl (20 mg/ml) in pyridine, sealed, and each sample was incubated with 20 μl of BSTFA for 1 hour before GC-MS analysis. In brief, derivatized samples (1–2 μl) were injected in split or split-less mode onto a DB-5MS capillary column (30 m × 0.25 mm × 0.25 μm; Agilent Technologies) using an Agilent 7890B gas chromatograph coupled to a 5977A mass selective detector. Helium was used as the carrier gas at a constant flow rate of 1 ml/min. The GC oven was operated with the following temperature program: initial temperature 70 °C (held for 2 min), ramped to 300 °C at 10 °C/min, and held at 300 °C for 5 min. The injector temperature was maintained at 250 °C, and the transfer line was set to 280 °C. Mass spectra were acquired under electron ionization (EI) at 70 eV, scanning from m/z 50–650 at a rate of ∼2 scans/s. Quality control samples and calibration standards were run intermittently throughout the batch to ensure retention time stability and instrument performance. Metabolite identification was performed by matching mass spectra and retention indices against NIST library, followed by peak integration and quantification using Agilent MassHunter software.

## Results

### Expression and subcellular localisation of *Pf*AQP-GFP fusion protein in transgenic parasites

A sequence search in PlasmoDB identified a putative *Plasmodium falciparum* Aquaglyceroporin (*Pf*AQP), encoding a 258-amino acid protein with an Aquaporin-like domain (8-243aa, Pfam Accession No. PF00230, E-value: 4.3e^-43^) and six predicted transmembrane domains (Figure 1A). To elucidate the evolutionary relationship of *Pf*AQP, we constructed a maximum likelihood phylogenetic tree *Pf*AQP clustered tightly with other *Plasmodium* AQPs, forming a well-supported clade distinct from both human AQPs and bacterial GlpF (Figure S1A). To further characterize *Pf*AQP, we performed multiple sequence alignment with *E. coli* GlpF, *Hs*AQP9, and *Hs*AQP10 (Figure S1B). *Pf*AQP contains the hallmark Asn–Pro–Ala (NPA) motifs, which are defining feature of all aquaporins. Additionally, *Pf*AQP harbours a conserved aromatic/arginine (ar/R) selectivity filter, although with subtle amino acid substitutions compared to human AQPs. These variations are predicted to fine-tune solute selectivity, consistent with the role of *Pf*AQP as an aquaglyceroporin capable of transporting both water and glycerol. Topology analysis showed that *Pf*AQP possesses the typical aquaporin architecture with six transmembrane helices and cytoplasmic N- and C-termini (Figure S1B). The conserved NPA motifs in loops B and E converge within the pore to form the solute selectivity barrier, and its overall structure closely resembles that of GlpF and HsAQP9, indicating strong structural conservation across species. Overall, *Pf*AQP is a conserved aquaglyceroporin adapted for dual water and glycerol transport in *Plasmodium*. To understand the putative role and to study localization of *Pf*AQP, transgenic parasite line expressing *Pf*AQP-GFP fusion protein was generated. Western blot analysis of total parasite lysates using anti-GFP antibody detected a prominent band at ∼55 kDa, corresponding to the full-length *Pf*AQP-GFP (Figure 1B), confirming the expression of the fusion protein. Live-cell fluorescence and confocal microscopy revealed that *Pf*AQP-GFP was predominantly localized at the parasite periphery during trophozoites and early schizonts, suggesting membrane association (Figure 1C). In mature schizonts, *Pf*AQP-GFP exhibited a distinct honeycomb-like pattern, characteristic of the parasite plasma membrane (PPM) surrounding individual merozoites. In addition, some of the parasites at trophozoite stage also showed fluorescence in close proximity and around the food-vacuole (Figure 1D). Co-staining with the lipid membrane probe BODIPY-TR ceramide further validated these observations; BODIPY labelling showed partial co-localization with *Pf*AQP-GFP signals both parasite membrane and the food vacuole membrane, confirming its association with multiple membrane systems in the parasite. Indeed, we have recently showed that *Pf*AQP associates with purified food vacuole membranes. To further assess its localization within the parasitophorous vacuole (PV), immunolabelling of transgenic parasites was done using anti-SERA5, a well-characterized PV-resident protein. In schizont stages, SERA5 localizes to the PV space surrounding clusters of merozoites, while *Pf*AQP-GFP remained associated with the membranes of individual merozoites (Figure 1E), with no significant overlap. This clearly supports that *Pf*AQP is predominantly localized to the parasite membrane and the food vacuole membrane.

**Figure 1:**
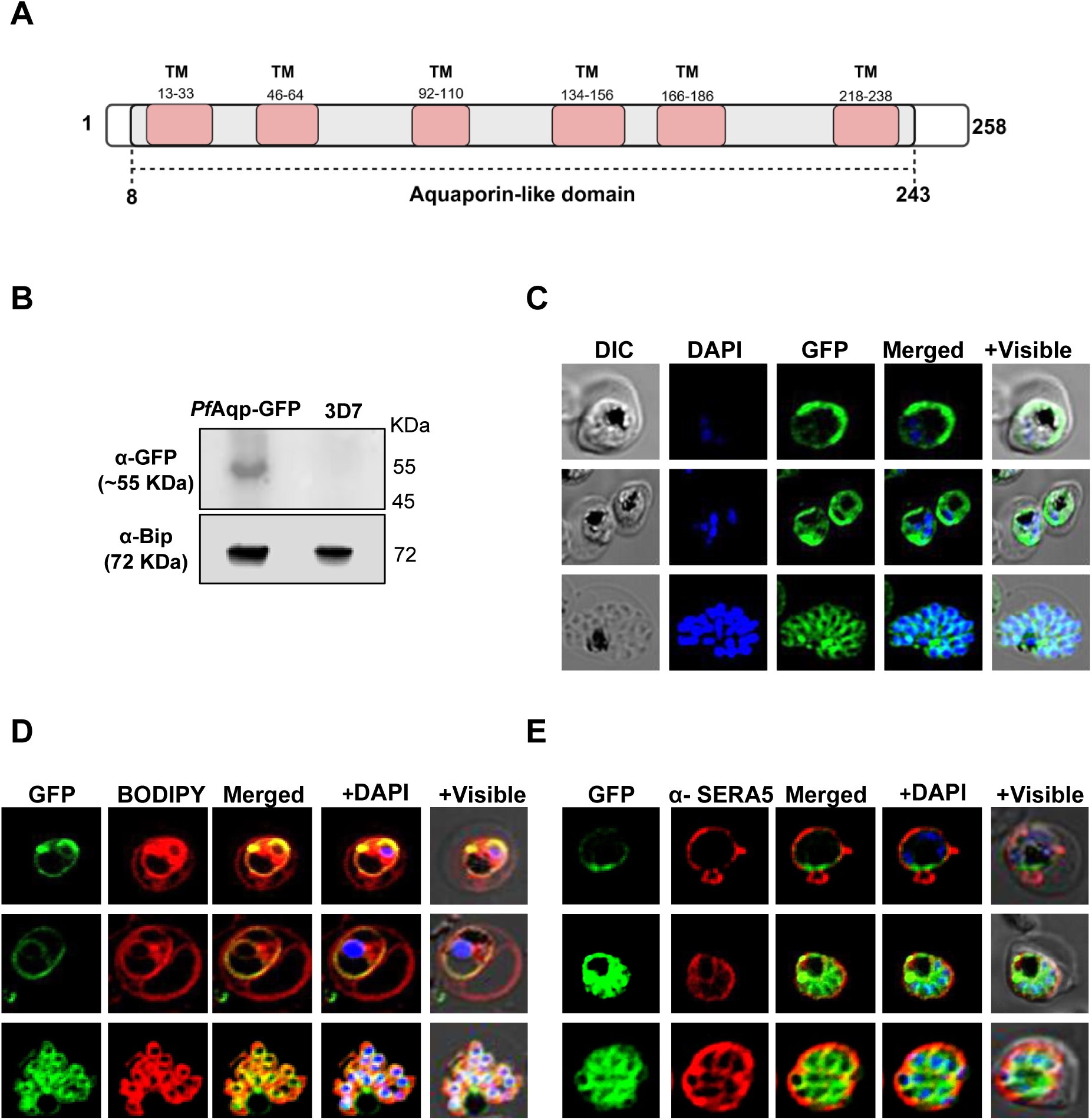
Expression and localization of *Pf*AQP-GFP fusion protein in transgenic parasites. A. Schematic showing domain organization of *Pf*AQP that harbors six transmembrane domains Aquaporin like domain (8-243 aa). Six transmembrane domains (TM) were denoted with respective amino acid positions. B. Western blot analysis of lysate of transgenic and wild-type parasites using an anti-GFP antibody. The fusion protein band (∼55 kDa) was detected in the transgenic parasites only (lane 1) and not in the wild-type parasites (lane 2). Blot ran in parallel with an equal amount of the same sample, probed with anti-BiP antibody was used as a loading control. C. Expression and localization of *Pf*AQP-GFP fusion protein through asexual parasite developmental stages; the parasite nuclei were stained by DAPI (blue) and parasites were visualized by a confocal laser scanning microscope. *Pf*AQP-GFP fusion protein fluorescence was detected around the parasite periphery as well as associated with the food-vacuole membrane. In +Visible panel, scale bar = 2 μm. D. Fluorescent microscopy images of transgenic parasites expressing *Pf*AQP-GFP fusion protein, stained with membrane probe BODIPY™ TR ceramide, showing fluorescence of GFP fusion protein overlapping with the membrane labeling at the parasite periphery (PPM/PVM) as well as at the food-vacuole boundary. Parasite nuclei were stained with DAPI and images were acquired by confocal laser scanning microscope. In +Visible panel, scale bar = 2 μm. E. Fluorescent microscopy images of transgenic parasites immuno-stained with anti-SERA5 antibody. GFP fluorescence at the parasite boundary did not show clear overlapping with anti-SERA5 staining; the GFP fluorescence showed clear honeycomb like structure in schizont stages (panel 2 and 3), whereas SERA5 staining was seen majorly in the PV region. Parasite nuclei were stained with DAPI and images were acquired by confocal laser scanning microscope. Scale bar = 2 μm.

### Endogenous tagging of *pfaqp* gene for transient knock-down in transgenic parasites

To investigate the functional role of *Pf*AQP, an inducible knockdown parasite line was generated by integrating a C-terminal HA-*glm*S ribozyme tag into the endogenous *pfaqp* locus using the selection-linked integration (SLI) strategy (Figure 2A). This approach enabled expression of the *Pf*AQP-HA fusion protein under its native promoter. The incorporation of a C-terminal tag in the native *pfaqp* gene was confirmed by diagnostic PCR-based analysis using genomic DNA from clonally selected parasite populations (Figure 2B). The integrant parasites showed expression of *Pf*AQP-HA fusion protein of ∼30 kDa, the expected size for the endogenously HA-tagged protein (Figure 2C). Such a band could not be detected in the parental 3D7 parasite line. The expression of the fusion protein was also confirmed by immunofluorescence assay (Figure 2D). *Pf*AQP-HA fusion protein showed similar localization pattern and association with the parasite membrane (Figure S2), as shown by localization studies with *Pf*AQP-GFP fusion protein.

**Figure 2:**
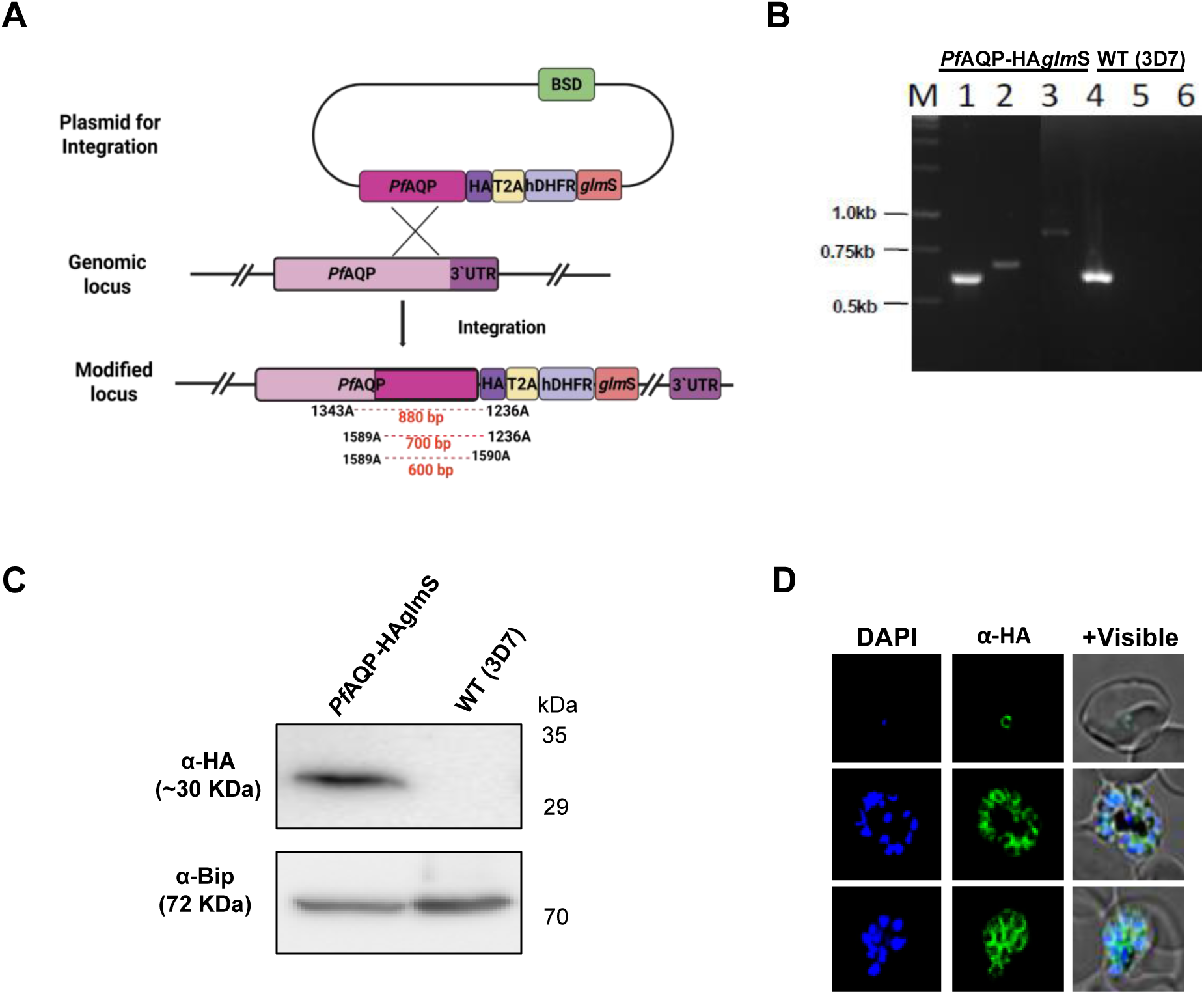
*Pf*AQP is essential in asexual replication and stage transition of *P. falciparum*. A. Schematic representation illustrating the single cross-over homologous recombination resulting in the incorporation of HA-*glm*S tag at the C-terminus of endogenous *Pf*aqp gene. B. PCR-based analyses to confirm integration of plasmid in the target gene locus using total DNA of the *Pf*AQP-HAglmS transgenic parasite culture and wild type WT (3D7) parasite lines, locations of primers are marked in the schematic (Fig. 3A). Lane 1 and 4: amplified using primer 1589A and 1590A, showing amplification of C-terminal fragment in transgenic parasites with desired integration as well as in parental 3D7 parasite line; lane 2 and 5: amplified using primer 1589A and 1236A, showing amplification of C-terminal along with HA tag in integrants or episomal parasites; lane 3 and 6: amplified using primer 1343A and 1236A, showing amplification of full length *pfaqp* gene along with HA tag in the integrants only. C. Western blot analysis of lysate of transgenic and wild-type parasites using anti-HA antibody. The fusion protein band (∼30 KDa) was detected in the transgenic parasites, *Pf*AQP-HAglmS line (lane 1) and not in the 3D7 parent parasite line (lane 2). Blot ran in parallel with an equal amount of the same sample, probed with anti-BiP antibody was used as a loading control. D. Fluorescent microscopic images of transgenic parasites, immunolabelled with anti-HA antibody showing expression of the *Pf*AQP-HA fusion protein throughout the asexual stage of the parasite. Parasite nuclei were stained with DAPI and images were acquired by confocal laser scanning microscope. In +Visible panel, scale bar = 2 μm.

### Inducible knock-down (iKD) of *Pf*AQP retards parasite-growth and hinders asexual cell-cycle

The transient knockdown of *Pf*AQP in transgenic parasites cultures grown with GlcN (2.5mM and 3.5 mM) or solvent alone (control) were analysed by western blot analysis (Figure 3A). The parasites in treated set (*Pf*AQP-iKD) clearly showed ∼80% reduction in *Pf*AQP levels as compared to control. To evaluate the functional significance of *Pf*AQP during asexual blood stage development, parasitemia was monitored in (*Pf*AQP-iKD) and control set, over two intraerythrocytic cycles (at 48 h and 96 h). Glucosamine-induced knockdown of *Pf*AQP led to a marked reduction in parasite replication, ∼84% decrease in parasitemia was observed in the *Pf*AQP-iKD set compared to control by 96 h, indicating that *Pf*AQP is essential for efficient asexual stage proliferation (Figure 3B). Furthermore, there was no deleterious effect of 3.5 mM GlcN on the growth of the wild-type parasite line (Figure S3). To assess the impact of *Pf*AQP depletion on parasite development, we examined the morphology and stage progression of *Pf*AQP-iKD and control parasites at defined time points during the asexual cycle. In control sets, parasites progressed normally from rings to trophozoites and mature schizonts, followed by efficient merozoite release and invasion, resulting in a 5–6-fold increase in parasitemia per cycle. In contrast, *Pf*AQP-iKD parasites exhibited normal morphology during ring stages, but showed clear developmental defects at the trophozoite stage, which appeared shrunken and poorly developed. Progression to mature schizonts was severely impaired, with numerous abnormal schizonts displaying incomplete nuclear division as well as merozoite development; indeed, these abnormal schizont lack distinct daughter merozoites. Stage composition analysis confirmed a developmental block from trophozoite to schizont stages in the knock-down set, leading to a marked reduction in new ring-stage formation in the next cycle (Figure 3C-D).

**Figure 3:**
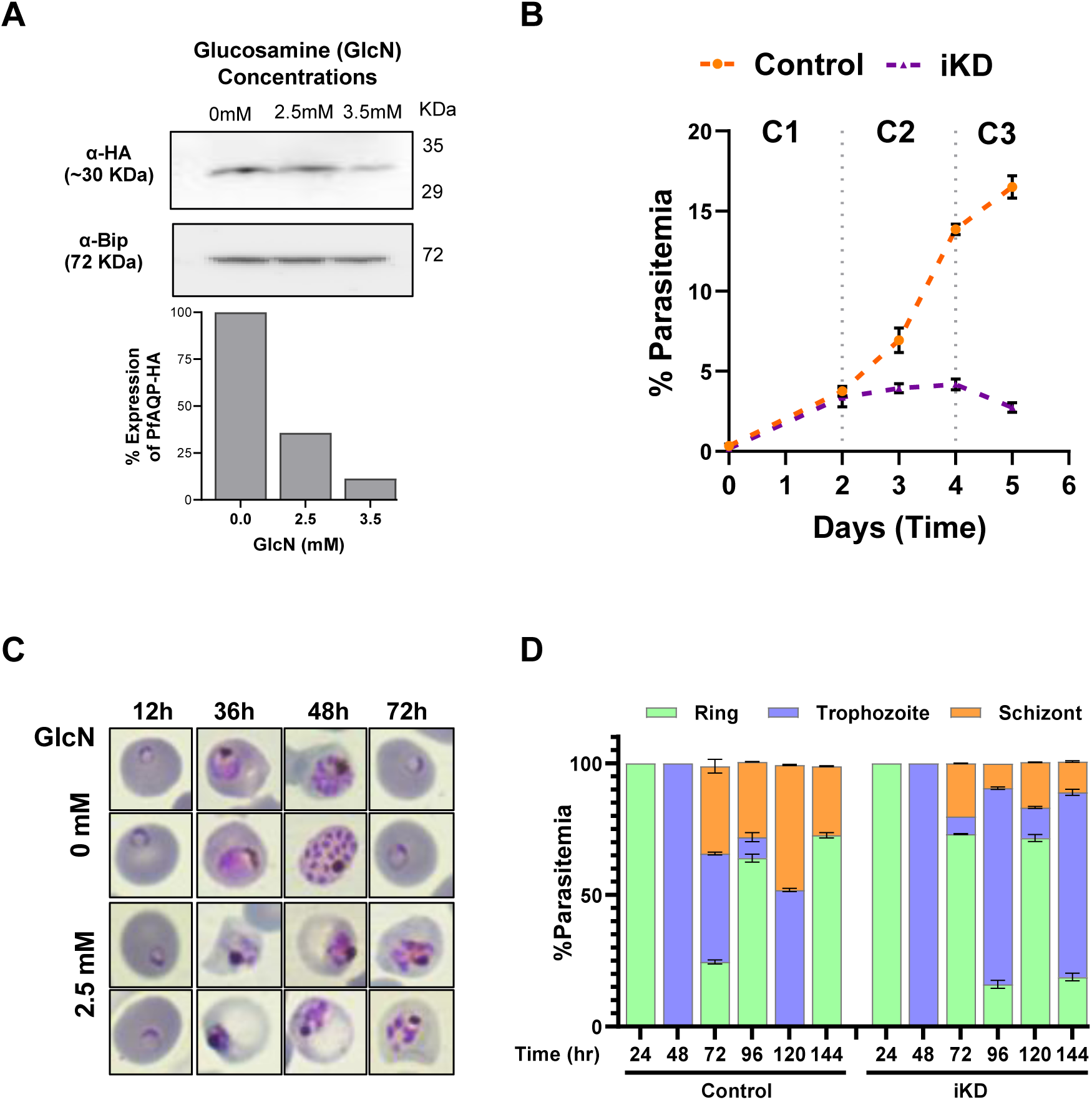
*Pf*AQP is essential in asexual replication and stage transition of *P. falciparum*. A. Immunoblot analysis of transgenic parasite lysate grown with different concentration glucosamine (0mM, 2.5mM and 3.5mM), probed with anti-HA antibody, showing a reduction in the fusion protein levels in presence of glucosamine. A blot ran in parallel with the same samples and probed with an anti-BiP antibody is used as the loading control. In bottom panel, graph showing the densitometry analysis of western blot results, indicating the expression levels of *Pf*AQP-HA fusion protein in response to treatment with different concentrations of GlcN or the solvent control (0mM). The data was normalized using the expression levels of BiP. B. Graph showing growth of transgenic parasites in presence of glucosamine (-iKD) in comparison to untreated control, over three asexual cycles; growth was determined by counting total parasitemia at 48, 96, and 144 hpi. All analyses were carried out in triplicate and error bars represent the standard deviation. C. Stage distributions of parasite from control and *Pf*AQP-iKD sets. Left panel: representative images of Giemsa-stained blood films at described time points (12, 36, 48, and 72 h). D. Parasite stage composition of transgenic parasites grown without or with glucosamine (control and *Pf*AQP-iKD sets respectively) at different time points during the intraerythrocytic cycle.

### Transient knock-down of *Pf*AQP disrupts parasite lipid homeostasis

The aquaporins are shown to associate with lipid biogenesis in eukaryotic cell, and specifically effecting the synthesis of triglycerides, through regulating the glycerol uptake ^16^. We have earlier shown in *P. falciparum* that TAGs and DAGs stores regulates the timely synthesis of PLs, which are essentially required for biogenesis of membrane, and hence for the parasite development as well as for merozoites formation during schizogony ^4^. Therefore, we assessed any role of the *Pf*AQP in parasite lipid homeostasis, its putative associated function with phospholipid synthesis and membrane biogenesis. First the changes in FA content were analysed by GCMS approach, which revealed a significant reduction in total FA content under *Pf*AQP-iKD conditions (Figure 4A). Detailed analysis of the FA composition of total FA revealed significant reductions in the abundance of C16:0, and C18:1(cis), all of which are known to be massively scavenged from the host (Figure 4B, S4). Further to determine which lipid species were affected by the loss of *Pf*AQP, untargeted lipidomic profiling of total lipid extracts was analysed using Independent Data Acquisition (IDA) strategy. The amount of variation in lipidome profile between control and iKD data sets were assessed using principal component analysis (PCA). Differential lipid analysis between control and *Pf*AQP-iKD parasites revealed a striking remodelling of the lipid landscape, and the score plot of the overall lipid abundance showed clear segregation between control and iKD sets for total lipids, with a variance of PC1 84.7% and PC2 4.1%, as well as for phospholipids, with a variance of 84.5% and PC2 4.5% (Figure S5). Several lipid species were significantly downregulated upon *Pf*AQP depletion, including a broad range of phosphoinositides (PIP, PIP2, PIP3) and ceramide derivatives (Cer, CerCl), suggesting disruptions in membrane signalling and trafficking pathways. In contrast, the knockdown led to accumulation of different species of monoacylglycerols (MAGs), triacylglycerols (TAGs), phosphatidic acid (PA), CDP-diacylglycerol (CDP-DAG), sphingomyelin (SM), and free fatty acids (FFAs), pointing to increased storage lipid biosynthesis (Figure 4C). To specifically assess the effect of *Pf*AQP knockdown on phospholipid composition, a focused analysis of the phospholipid subset from the total lipidome was carried out using IDA. Several phosphatidylinositol phosphates (PIPs), including PIP2 37:6 and PIP2 38:4, along with lysophosphatidic acid (LPA 17:2), phosphatidic acids (PA), phosphatidylinositols (PI), and CDP-diacylglycerol (CDP-DAG 33:3), were significantly downregulated in the knockdown condition. Conversely, numerous phospholipid species were markedly upregulated, including lysophosphatidylinositol (LPIP), phosphatidylinositol phosphates (PIP, PIP2, PIP3), and N-acyl-phosphatidylethanolamine (NAPE 42:5+NH4), indicating enhanced phospholipid turnover or compensatory membrane remodelling (Figure 4C, S6).

**Figure 4:**
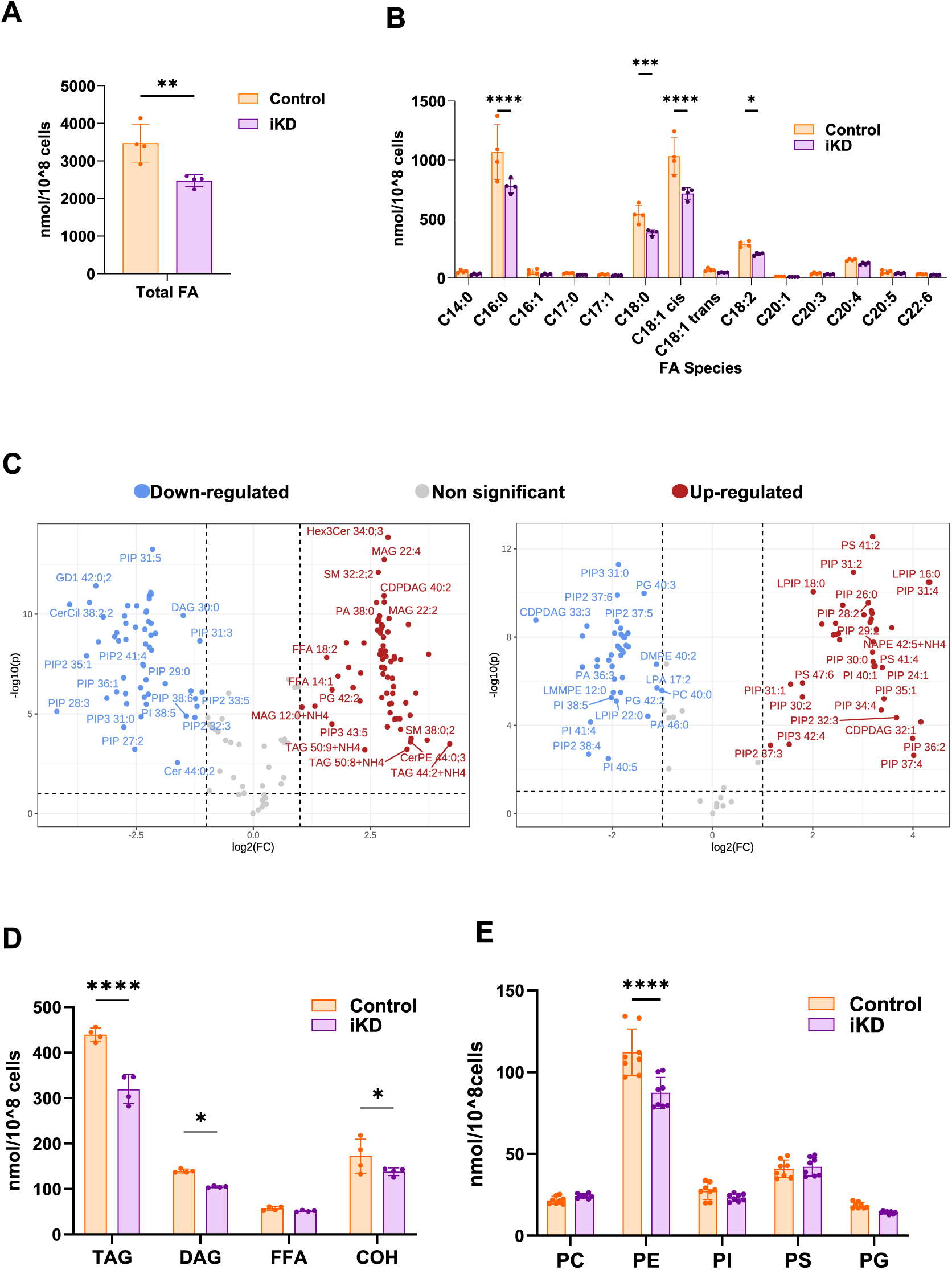
Ablation of *Pf*AQP results in lipid homeostasis disruption and reduction in the level of major phospholipids in the parasite. A. Bar graph showing total fatty acid abundance (in nmol lipid/10^7parasites) in control and *Pf*AQP-iKD sets, as determined by mass spectrometry-based lipidomic analyses. B. Analysis of fatty acids (FAs) showing no significant changes in the FA composition of parasites from control and *Pf*AQP-iKD sets. C. Volcano diagrams showing the changes global lipidome profile in total lipids (left panel) and phospholipids (right panel) of control vs *Pf*AQP-iKD sets acquired through Independent Data Acquisition (IDA). The red dot on the right side of the figure represents the upregulated lipid species, the blue dot on the left side represents the downregulated lipid species and grey dots represent non-significant difference. The x-axis corresponds to log2 (fold change), and the y-axis corresponds to −log10 (p-value), the dots farthest from the origin ‘0’ represent the relatively higher level of significance. D. Analysis of major neutral lipids levels showing upregulation triacylglycerols (TAGs) of and downregulation of diacylglycerols (DAGs), cholesterols (CHO) in *Pf*AQP-iKD set. E. Analysis of the parasite’s phospholipid composition of different phospholipids of control and *Pf*AQP-iKD sets. Significant reduction in the level of PE was observed in *Pf*AQP ablated condition in comparison to the control, while the rest of the PLs remain unaltered. All analyses were performed in triplicate (n = 3) or more; the error bars indicate standard deviations. **** signifies p<0.0001, while *** signifies p<0.001, ** signifies p<0.01 and * signifies p<0.05. The p values were determined by unpaired Student’s *t*-test. PS, phosphatidylserine; PI, phosphatidylinositol; PC, phosphatidylcholine; PG, phosphatidylglycerol; PE, phosphatidylethanolamine; SM, sphingomyelin; PL, phospholipid.

### Knockdown of *Pf*AQP disrupts synthesis of neutral lipids and membrane phospholipids

Given the function of aquaglyceroporins, the knockdown of aquaglyceroporin is expected to limit glycerol availability, suppressing glycerol-3-phosphate production and downstream DAG synthesis. This can create a bottleneck in neutral lipid and phospholipid biosynthesis, especially *de novo* synthesis of PC and PE, through Kennedy pathway, and associated synthesis of PS. To assess effect of *Pf*AQP-iKD on biosynthesis of membrane phospholipids and neutral storage lipids, comparative lipidomics analysis between the control and knockdown (iKD) parasite lines were carried out. The abundance of neutral lipids, and PLs were found to remained unchanged in the absence of *Pf*AQP (Figure S7A), however there was a significant reduction in diacylglycerol (DAG) and triacylglycerol (TAGs) levels in the *Pf*AQP-iKD line (Figure 4D, S8), which is indicative of impaired de novo glycerolipid synthesis. Additionally, cholesterol level was also significantly reduced in the iKD condition (Figure 4D,S8), possibly reflecting decreased membrane synthesis demand or reduced uptake from the host environment. Further, the phospholipid composition revealed significant changes in *Pf*AQP-iKD set. Among the major phospholipids, phosphatidylethanolamine (PE) exhibited a significant decrease in content upon *Pf*AQP depletion, indicating a disruption in PE biosynthesis pathways (Figure 4E and S7B). Interestingly, phosphatidylserine (PS) levels remained largely unchanged or showed a modest increase (in Mol%), suggesting stable or reduced turnover rather than *de novo* synthesis alteration. The phosphatidylcholine (PC) levels also remain unchanged or showed minor increase (in Mol%) in the knockdown set as compared to controls (Figure 4E, S7B), which may suggest compensatory upregulation via scavenging (lysoPC) or enhanced choline pathway despite impaired DAG supply. Detailed analysis of FA composition of PE molecular species revealed significantly altered FA abundance, homeostasis and combinations: all PE species containing short, medium, to long FA species, i.e. molecular species containing combination of C14:0, C16:0, were found significantly reduced (e.g.: C34:1/2), whereas all PE species containing very medium or long FA combinations such as C36:4 were significantly increased (Figure S9). Importantly, further analyses of the FA compositions of the two phospholipid species, PC and PS, also revealed significant alteration in their FA content in PfAQP-iKD set in comparison to control. In contrast, in the subset of other phospholipids, phosphatidylinositol (PI) and phosphatidylglycerol (PG) both showed moderate decrease in the knockdown condition (Figure 4E, S7B). Collectively, these results suggest that *Pf*AQP plays a pivotal role in regulating lipid homeostasis and membrane biogenesis by modulating availability of key components for phospholipid biosynthesis.

### *Pf*AQP knock-down parasites show disrupted/underdeveloped plasma membrane

*Pf*AQP-deficient parasites failed to form proper segmented schizonts, along with disrupted lipid homeostasis. To further elucidate the severity of *Pf*AQP knock-down on segmentation defects and membrane biogenesis, ultrastructure expansion microscopy (U-ExM) was conducted on synchronized *P. falciparum* schizonts stage parasites from control and *Pf*AQP depletion conditions. Plasma membrane staining of parasites with anti-*Pf*MSP1 antibody showed dysmorphic plasma membrane staining with aggregates of incompletely segmented merozoites upon the *Pf*AQP knockdown (Figure 5, S10). In more details, in the control parasites distinct membrane staining with *Pf*MSP1 was observed surrounding individual nuclei, indicating well-defined parasite membranes and proper segmentation of daughter merozoites. The nuclei appeared regularly distributed and discrete, consistent with normal schizont maturation, collectively represent intact membrane biogenesis and organized nuclear division (Figure 5, upper panel). In contrast, *Pf*AQP-depleted parasites exhibited striking structural abnormalities. The *Pf*MSP1 signal was diffuse and discontinuous, suggesting severe impairment in membrane formation and stability. Many schizonts displayed collapsed or fragmented membrane structures, accompanied by irregular or clustered nuclear staining, along with compromised schizont segmentation and defective organellar partitioning (Figure 5, bottom panel). These abnormalities likely reflect disrupted glycerol transport and subsequent perturbation of lipid biosynthesis pathways, leading to defective phospholipid assembly and membrane integrity.

**Figure 5:**
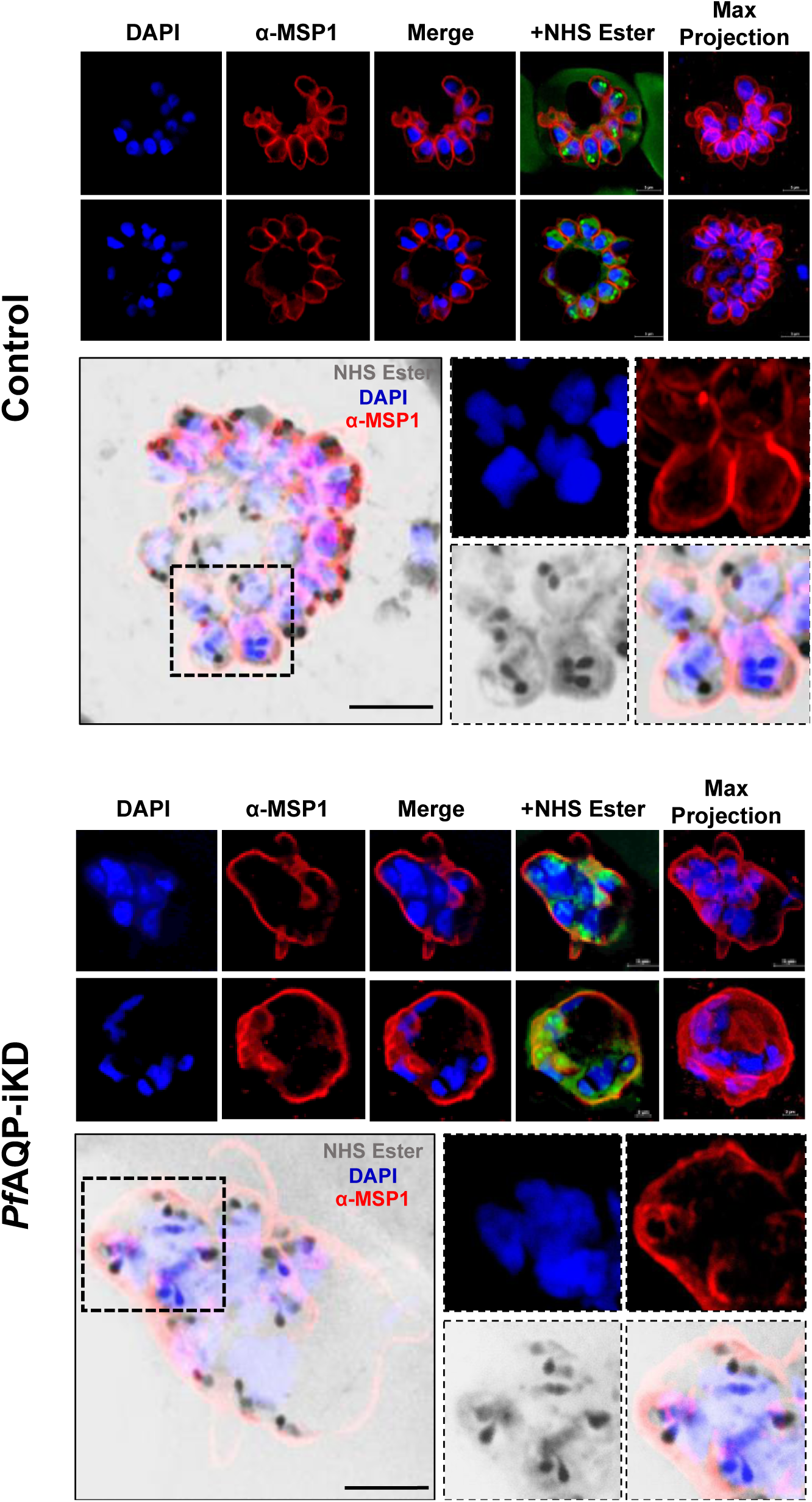
*Pf*AQP -deficient parasites show disrupted/underdeveloped plasma membrane. U-ExM of *Pf*AQP parasites comparing segregation and development of plasma membrane in *Pf*AQP control and iKD conditions. Late schizont-stage parasites were stained with anti-MSP1antibody to stain the parasites membrane marker in red. NHS Ester was used as a general protein stain and is shown in green (upper portion), grey (in zoomed one) and nuclei (DAPI) are in blue. Maximum projections of approximately 29 z-slices are shown on the right. To the bottom panel of each large image are zoomed-in views of selected parasite comparing segregation and development of plasma membrane in *Pf*AQP control and iKD conditions. For the zoomed-in images, the selected merozoites were selected from a subset of z-slices from the larger image. In +Visible panel, scale bar = 5 μm.

### Ablation of *Pf*AQP causes reduction in amino-acid levels in parasites and dysregulate mitochondrial energy metabolism

Reduction in cellular glycerol level can influence amino acid metabolism for gluconeogenesis, in addition disrupt the generation of neutral lipid species ^35, 36, 37^. Therefore, targeted quantification of intracellular amino acid levels in *Pf*AQP-iKD parasites was carried out; knock-down parasites exhibited a substantial reduction in several amino acids. Notably, among the amino acids obtained primarily or exclusively from haemoglobin, leucine, serine, threonine and tyrosine showed significant reduction; similarly, among the amino acids obtained exclusively or primarily from host milieu directly, isoleucine, methionine and glutamic acid showed significant reduction. In contrast, glycine, proline, and arginine levels remained relatively unchanged (Figure 6A). The broad downregulation of amino acid pools in *Pf*AQP-depleted parasites suggests metabolic rewiring likely triggered by altered nutrient flux.

**Figure 6:**
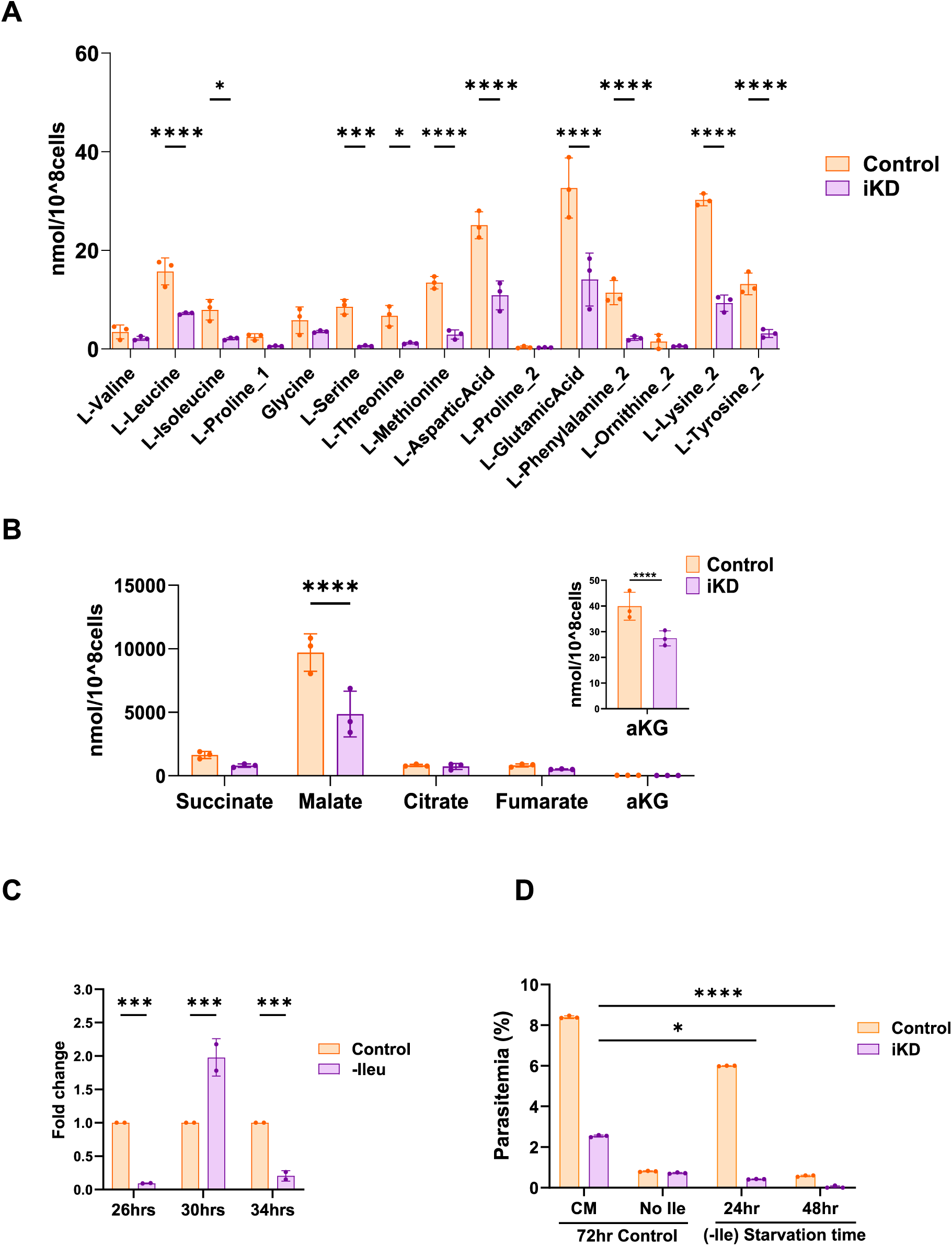
*Pf*AQP depletion perturbs the amino acid metabolism and TCA cycle. A. Targeted metabolomics analysis of control and *Pf*AQP-iKD parasite cultures, using liquid chromatography-mass spectrometry (LC-MS). Bar graph representing the concentrations of selected amino acids in control and *Pf*AQP-iKD parasite, showing that under *Pf*AQP-iKD conditions, there was significant reduction for amino acids including glutamic acid, aspartic acid, leucine, isoleucine and serine. Data are shown as mean ± SD (*n* = 3), statistical significance is indicated by \**p* < 0.05 compared to baseline levels. B. Targeted metabolomics analysis of control and *Pf*AQP-iKD parasite cultures, using liquid chromatography-mass spectrometry (LC-MS). Bar graph showing fold changes/levels (nmoles/10^8 cells) of TCA cycle metabolites in the control and CLS-iKD sets. Each bar represents the mean concentration ± SD (*n* = 3 biological replicates) for key TCA cycle intermediates: citrate, α-ketoglutarate, succinate, fumarate and malate.

The glycerol uptake and utilization also associate with carbon metabolism, in addition to lipid and amino acid metabolism; further, the incomplete/branched TCA cycle of the parasite specifically depends upon conversion of glutamate to α-ketoglutarate, which subsequently gets incorporated in the cycle (^7, 38, 39^). In addition, *Pf*AQP-iKD caused reduction of amino-acids levels in the parasite including the glutamate levels. Therefore, impact of *Pf*AQP depletion was also assessed on central carbon metabolism in the parasite. Targeted comparative metabolomic profiling for tricarboxylic acid (TCA) cycle intermediates in control and *Pf*AQP-iKD sets revealed that malate and α-ketoglutarate levels were markedly reduced in the *Pf*AQP-iKD condition. In contrast, levels of citrate were nearly unchanged (Figure 6B). This pattern suggests that *Pf*AQP knockdown leads to a dysregulated TCA cycle, potentially due to impaired metabolite exchange. These findings highlight a possible link between *Pf*AQP function and mitochondrial metabolic balance in the malaria parasite.

### Role of *Pf*AQP in maintenance of survival strategies under starvation induced stress

It has been shown that starvation and fasting facilitates enhanced glycerol uptake in eukaryotic cells ^40^; further, our data showed that *Pf*AQP-iKD disrupts uptake of crucial amino acids by the parasite from host milieu. Therefore, the possible role of *Pf*AQP to support parasite survival under amino acid starvation conditions was assessed. To assess any change in the expression levels of *Pf*AQP in parasites growing in the isoleucine-free media, the relative mRNA levels of *pfaqp* in complete media or isoleucine free media (-Ile) were determined. During the initial starvation period (at 26hpi), parasites showed significant reduction in relative mRNA levels of *pfaqp*. However, when parasites were starved till the trophozoite stage (at 30 hpi), when hibernation stage is initiated ^41, 42, 43^, there was >2-fold increased expression levels of *pfaqp*. This increase in expression remained only for brief period; parasites starved for longer period (>30 hr), showed decreased expression levels of *pfaqp* (Figure 7A).

**Figure 7:**
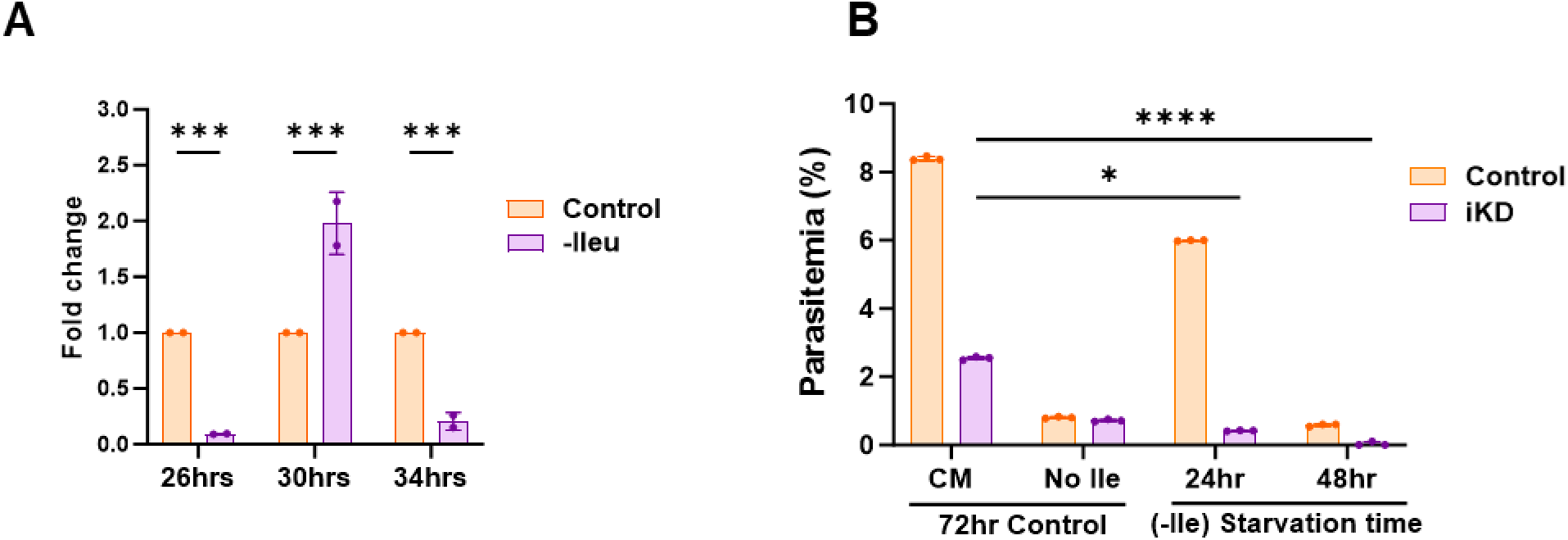
*Pf*AQP plays role in parasite survival under starvation induced stress. A. Quantitative RT-PCR showing relative *pfaqp* expression in *P. falciparum* 3D7 parasites at different hpi after growth in Ile-free media versus complete media, demonstrating *Pf*AQP upregulation during the hibernatory stage under amino acid starvation. B. Graph depicting recovery of parasite growth following isoleucine re-supplementation in control and *Pf*AQP-iKD parasites after 24 h or 48 h growth in Ile-free media. Parasitemia was measured after 72 h recovery in complete media, with parasites maintained continuously in complete (+Ile) or Ile-free (−Ile) media serving as positive and negative controls, respectively, demonstrating that *Pf*AQP ablation during isoleucine starvation impairs recovery.

To ascertain the role of *Pf*AQP in maintaining cellular homeostasis during amino-acid starvation, revival of parasite cultures after removal of starvation was assessed under *Pf*AQP knock-down conditions. Parasite cultures in control and *Pf*AQP-iKD set were grown in isoleucine free media for different time period, and subsequently supplemented with complete media to allow recovery. In control set, about 80% parasite recovered after 24hr of starvation, whereas only ∼10% parasites could recover after 48hr of starvation. In *Pf*AQP-iKD set only <5% of parasites could recover after 24hr of starvation, whereas almost no parasite was able to recover after 48hr of starvation. It is to be noted that ∼25% parasite survived at 72hpi time point in *Pf*AQP-iKD set grown in complete media, which suggest ∼75% reduction in revival efficiency for *Pf*AQP-iKD parasites (Figure 7B). Overall, these results show the role of *Pf*AQP in cell survival when it undergoes cellular stress.

## Discussion

Efficient import of nutrients and export of metabolic waste across membrane systems is fundamental for the survival and development of *Plasmodium* parasites throughout their complex life cycle. This exchange is mediated by a repertoire of transmembrane transporters localized across various membranes including the parasitophorous vacuole membrane (PVM), plasma membrane, and internal organellar membranes. These transporters have been established as critical determinants of parasite viability and pathogenicity ^44^. Among them, aquaporins-water channel proteins conserved across nearly all living organisms-facilitate the rapid movement of water and small solutes, such as glycerol, to maintain cellular homeostasis. A subclass of these proteins, known as aquaglyceroporins, is permeable to both water and glycerol and has been identified in various *Plasmodium* species.

Here, we have deciphered functional role of homologue of Aquaglyceroporins (AQP) in the human malaria parasite *P. falciparum* (*Pf*AQP) through detailed localization, gene knock-down, and cellular functional analyses during the asexual blood stages of the parasite. To understand the potential role of *Pf*AQP in the parasite, we carried out detailed localisation by confocal microscopy using a transgenic parasite line expressing PfAQP-GFP fusion protein. Our results with transgenic parasite *Pf*AQP associate with the parasite plasma membrane as well as with the food-vacuole membrane. Our recent studies on proteomic analysis of isolated food-vacuoles also identified *Pf*AQP as a membrane associated protein ^45^.

To dissect the functional significance of *Pf*AQP in *Plasmodium falciparum*, we employed the *glmS* ribozyme-mediated inducible knockdown system. Selective downregulation of *Pf*AQP markedly impaired parasite growth and survival during the asexual blood-stage, underscoring its essential role in intraerythrocytic development cycle. Morphological examination revealed that *Pf*AQP knockdown disrupts the developmental transition from trophozoite to schizont, suggesting a defect in the late stages of parasite maturation. This impaired progression is likely linked to compromised glycerol uptake, as previously demonstrated in *P. berghei* where *Pb*AQP-null parasites exhibited severely reduced glycerol acquisition linked with growth phenotype ^19^. Notably, expression studies in *Xenopus* oocytes have shown that *Pf*AQP is permeable to both water and glycerol, further supporting its role in serum glycerol uptake ^16^. The imported glycerol can be channelled toward glycerophospholipid and phospholipid biosynthesis or into glycolysis, both of which are essential for membrane biogenesis and energy metabolism during rapid parasite replication.

We have earlier shown that the parasite uses host acquired phospholipids to generate fatty acids which are timely mobilized for synthesis of phospholipids essentially required for membrane biogenesis at schizogony ^4^. In addition, under high lipid biosynthetic demand during schizogony, extracellular glycerol acquisition could be also critical for the parasite. Therefore, to gain deeper mechanistic insight into the functional role of *Pf*AQP in glycerophospholipid and phospholipid biosynthesis, we conducted lipidomic analysis on lipids extracted from transgenic parasites under both control and knockdown conditions. Under normal physiological conditions, glycerol imported through *Pf*AQP is phosphorylated to glycerol-3-phosphate (G3P), a critical precursor for diacylglycerol (DAG) synthesis. DAG, in turn, contributes to the biosynthesis of major phospholipids such as phosphatidylethanolamine (PE), phosphatidylcholine (PC), and phosphatidylserine (PS). Our lipidomic analysis revealed that there was significant change in the FA content and composition in parasites lacking *Pf*AQP, pointing towards reduced scavenging from the host. The reduction of PE content in the mutant could indicate, a defect in the Kennedy pathway which corresponds to generation of DAG due to impaired glycerol uptake. Interestingly, PS levels remained stable or slightly increased indicating PS synthesis may be partially supported through the utilization mitochondrial PS decarboxylation or could result in PE depletion and PS accumulation equilibrium (base exchange of polar head groups). PC content remained unchanged (or slightly increased), which may reflect a compensatory response through lysoPC scavenging or increased flux through the CDP-choline branch of the Kennedy pathway. Additionally, mitochondrial phospholipids such as phosphatidylglycerol (PG) and phosphatidylinositol (Pi) also showed a decreasing trend, suggesting that lipid imbalance may extend to mitochondrial compartments, potentially affecting organelle function. Disruption in phospholipid homeostasis and synthesis showed clear effect on membrane biogenesis on the developing parasites; the late-stage parasites showed developing merozoite however poorly formed plasma membranes.

Meanwhile, the accumulation of TAG and reduction in DAG highlight a redirection of lipid flux toward neutral lipid storage. This lipid overflow response is a well-known stress adaptation in *Plasmodium*, allowing the parasite to store excess fatty acids that cannot be used for membrane synthesis due to G3P/DAG limitation. Finally, the reduction in cholesterol could be a secondary consequence of reduced membrane biogenesis demand or altered lipid trafficking and homeostasis under AQP-limited conditions. Taken together, these findings suggest that *Pf*AQP knockdown induces a metabolic shift characterized by impaired glycerol backbone availability, disrupted membrane phospholipid biosynthesis, and compensatory lipid scavenging and storage.

In addition to disruption of neutral lipid generation, reduction in cellular glycerol level can also influence amino acid metabolism ^7, 38^. Targeted quantification of intracellular amino acid levels in *Pf*AQP-iKD parasites showed significant reduction in several amino acids obtained from haemoglobin and those exclusively or majorly scavenged from host. A broad downregulation of amino acid pool may suggests altered metabolic flux or poor scavenging from host; indeed, knock down parasites showed poor membrane development, which may affect scavenging of nutrient from host. Further, the amino acid poo could be also catabolized to enter into energy metabolism pathways. Asexual and gametocytes stages of parasite primarily depend on glycolysis for ATP synthesis and survival; the parasite also harbours all the enzymes required for TCA cycle ^7, 46^; however, the flux through acetyl CoA derived from glucose contribute >10% of the cycle. The major carbon flux in TCA cycle depends upon conversion of glutamate to α-ketoglutarate, which subsequently gets incorporated in the cycle, and hence the TCA cycle of the parasite is suggested to be the incomplete/branched ^7, 47^. Although the asexual stages of *P. falciparum* primarily rely on glycolysis for energy production, the maintenance of the TCA cycle and hence the mitochondrial respiratory chain is crucial for the reoxidation of inner membrane dehydrogenases-such as dihydroorotate dehydrogenase, which plays a key role in de novo pyrimidine biosynthesis ^48^. Targeted metabolomic analysis after *Pf*AQP ablation showed major reduction in amino acid pool including glutamate, which also caused significant reduction in generation of TCA metabolites including α-ketoglutarate, malate and succinate. This metabolic disruption resulted in cellular stress on the parasite, which also contributed in hindering the parasite growth.

These results show that *Pf*AQP ablation disrupts glycerol flux, which cause major changes in lipid homeostasis and defect in TCA cycle; the compensatory shift may not be able to fully replenish the lost glycerol-derived flux, thereby affecting parasite development and growth. Overall, the data suggest that *Pf*AQP is an important transporter that plays a crucial role in the parasite’s metabolism-facilitating glycerol-dependent phospholipid biosynthesis and indirectly supporting the TCA cycle by maintaining a balanced input of carbon substrates. *Pf*AQP knockdown triggers lipid remodeling and perturbs central carbon metabolism, underscoring its importance in maintaining metabolic flexibility during intraerythrocytic development.

Earlier, studies have demonstrated that starvation and fasting conditions facilitate glycerol transport, implicating aquaglyceroporins (AQPs) in promoting cell survival under nutrient-deprived and stress conditions ^40^. In *P. falciparum*, isoleucine is an essential amino acid which is exclusively acquired from host milieu; isoleucine starvation induces hibernation like conditions in the parasite, where the hibernating parasite can recover in complete media with varying efficiency up to 72h of starvation ^41^. We observed an upregulation of *Pf*AQP expression following amino acid starvation, coinciding with the time point when parasites begin transitioning into a hibernation-like dormant state ^41, 42, 43^. This suggests that *Pf*AQP is transcriptionally upregulated as part of the parasite’s adaptive response to stress, potentially to enhance glycerol uptake. The increased availability of glycerol may support lipid biosynthesis required for membrane maintenance or remodeling during dormancy. Parasites with *Pf*AQP knockdown exhibited significant ablation in recovery after starvation, minimal growth restored was restored in iKD set starved for 24 h and almost no recovery was observed for the iKD parasites starved for 48 or more. Therefore, the functional importance of *Pf*AQP in stress tolerance is ascertained by the impaired recovery of *Pf*AQP knockdown parasites following nutrient starvation. Consistent with previous reports ^49^, reduced glycerol uptake has been associated with decreased lipid synthesis and membrane biogenesis, which may hinder the induction of essential survival mechanism under cellular and nutritional stress including autophagy. Indeed, in similar lines our data showed reduced synthesis of phospholipid and membrane biogenesis in *Pf*AQP knockdown conditions. In summary, *Pf*AQP is a critical component of *P. falciparum* metabolism, enabling glycerol import to support lipid biosynthesis, membrane biogenesis and mitochondrial function (Figure 8). Its role becomes especially important during nutrient stress, where it helps maintain metabolic flexibility, redox balance, and developmental progression during the asexual blood stages.

**Figure 8:**
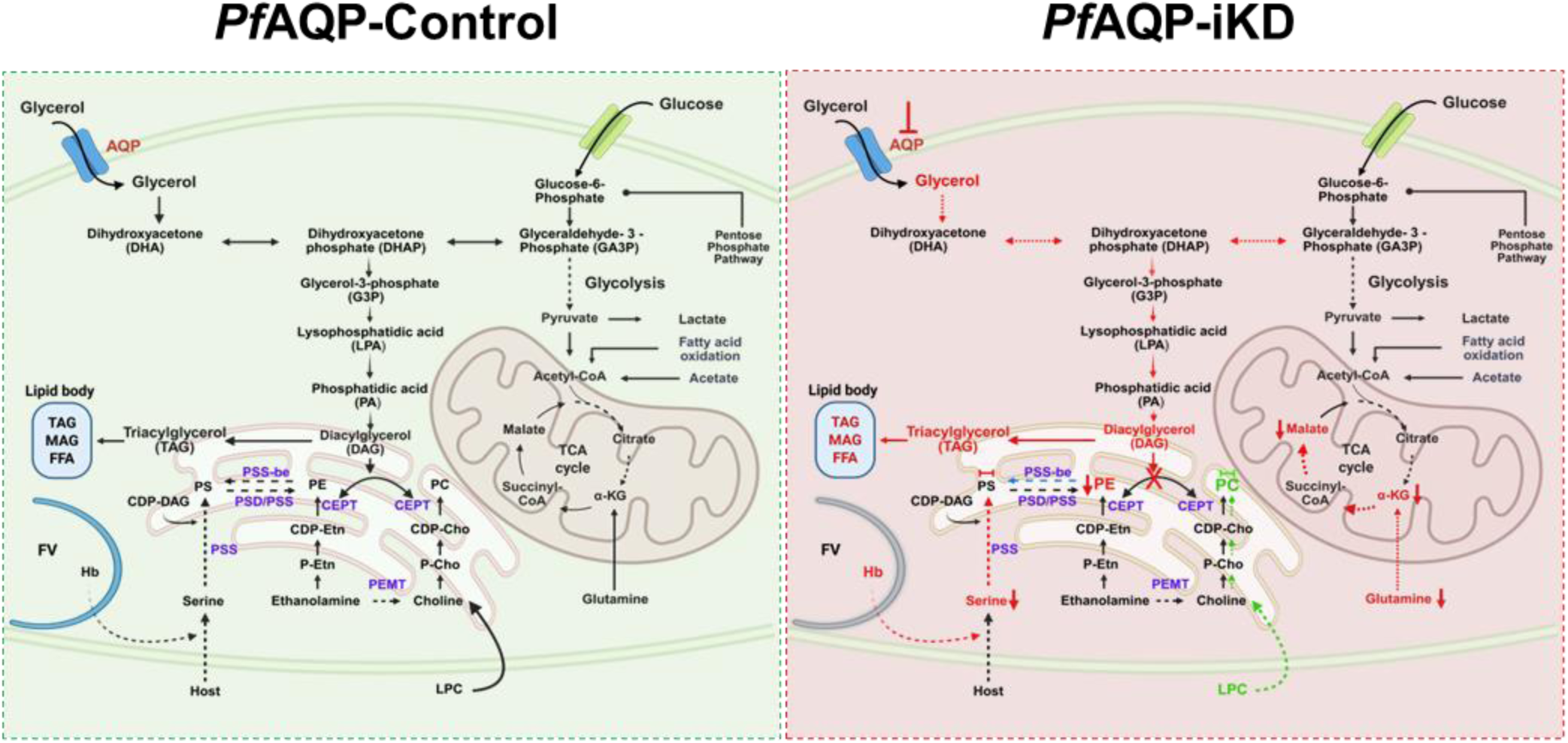
Schematic diagram showing the proposed role of *Pf*AQP in *Plasmodium falciparum* cell survival and lipid homeostasis. Under normal conditions, AQP mediates efficient glycerol uptake, supporting glycerol-3-phosphate (G3P) generation and downstream synthesis of LPA, PA, DAG, and TAG. Adequate DAG supply sustains de novo phospholipid biosynthesis, including PS and PE (via PSS, PSD/PSDS) and PC (via CEPT and PEMT), complemented by host-derived LPC salvage. Mitochondrial TCA cycle activity remains balanced, maintaining metabolic precursors and membrane biogenesis. Transient downregulation of *Pf*AQP, reduced glycerol imports limits G3P formation and constrains flux through the PA–DAG branch, resulting in diminished TAG storage and impaired synthesis of major phospholipids (PS, PE, PC). Perturbed lipid flux is accompanied by altered TCA cycle intermediates and increased dependence on host-derived serine and choline/LPC. Collectively, AQP depletion disrupts lipid homeostasis and compromises parasite viability.

## Data Availability

All relevant data are within the manuscript and its Supporting Information files.

## Competing interests

The authors declare that they have no competing interests

## Acknowledgements

We are grateful to Philip J. Shaw for providing vector pGFP_glmS; Siggi Sato for pSSPF2 vector. We thank Rotary blood bank, New Delhi for providing the RBCs. This work and was supported via a Collaborative Research Program Grant (Project 6003-1) and IARDP grant (#2023-0414) from Indo-French Centre for the Promotion of Advanced Research-CEFIPRA to CYB, YYB, and AM. The research work in AM’s and PM’s laboratories is supported Flagship Grant (#RAD-40/104/2023- MED-DBT) from the Department of Biotechnology, Govt. of India, Research grant (#CRG/2022/002655) from the Science and Engineering Research Board, Department of Science and Technology, (SERB-DST, Govt. of India), and Team Science Grant (IA/TSG/22/1/600422) from the DBT/Wellcome Trust India Alliance. CYB and YYB were supported by Agence Nationale de la Recherche, France (Project ApicoLipiAdapt grant ANR-21-CE44-0010; Project Apicolipidtraffic grant ANR-23-CE15-0009-01; Project OIL grant ANR-24-CE15-2171-02), The Fondation pour la Recherche Médicale (FRM EQU202103012700), Laboratoire d’Excellence Parafrap, France (grant ANR-11-LABX-0024), LIA-IRP CNRS Program (Apicolipid project), the Université Grenoble Alpes (IDEX ISP Apicolipid) and Région Auvergne Rhone-Alpes for the lipidomics analyses platform (Grant IRICE Project GEMELI). MI was supported by Pre-doctoral research fellowship from ICGEB. MN, MA were supported by research fellowships from the Department of Biotechnology, Govt. of India.

